# Characterization of adult human skeletal cells in different tissues reveals a CD90+CD34+ periosteal stem cell population

**DOI:** 10.1101/2022.12.05.519079

**Authors:** Ye Cao, Scott M. Bolam, Anna L. Boss, Helen C. Murray, Nicola Dalbeth, Anna E.S. Brooks, Brya G. Matthews

## Abstract

Skeletal stem and progenitor cells are critical for bone homeostasis and healing, but their identity and diversity in humans are not well understood. In this study, we compared stromal populations in matched tissues from the femoral head and neck of 21 human participants using spectral flow cytometry of freshly isolated cells. High-level analysis indicated significant differences in marker distribution between periosteum, articular cartilage, endosteum and bone marrow stromal populations, and identified populations that were highly enriched or unique to specific tissues. Periosteum-enriched markers included CD90 and CD34. Articular cartilage, which has very poor regenerative potential, showed enrichment of multiple markers, including the PDPN+CD73+CD164+ population previously reported to represent human skeletal stem cells. We further characterized periosteal populations by combining CD90 with other strongly expressed markers. CD90+CD34+ cells sorted directly from periosteum showed significant colony-forming unit fibroblasts (CFU-F) enrichment, rapid expansion, and consistent multi-lineage differentiation of clonal populations. In situ, CD90+CD34+ cells include a perivascular population in the outer layer of the periosteum and non-perivascular cells closer to the bone surface. In conclusion, our study indicates considerable diversity in the stromal cell populations in different tissue compartments within the adult human skeleton, and suggests that periosteal stem cells reside within the CD90+CD34+ population.

## Introduction

Tissue-resident skeletal stem and progenitor cells (SSPC) are critical for homeostasis and regeneration of the skeleton. Bone marrow is considered a fundamental niche for stem cells, but the periosteum plays a crucial role in bone healing. In contrast, articular cartilage has minimal tissue turnover and regeneration capacity. Previous studies concluded that perivascular mesenchymal stem cells that contribute to skeletal homeostasis are present in most tissues and have a common set of markers (1). However, more recently it has been recognized that there is incredible diversity in stromal populations even within a single tissue, and that different skeletal compartments probably have unique stem and progenitor cell populations (2-7). Therefore, it is unlikely that a single hierarchy of cells contributes to skeletal growth, homeostasis, and healing throughout the skeleton and markers for SSPC identification established using fetal or juvenile tissue, or in cell culture, may not be directly applicable to adult settings.

In mice, numerous markers and marker combinations are proposed to identify SSPCs (8). Some identify cells in the growth plate, a structure specifically involved in longitudinal growth that fuses in young adult humans (9, 10). Others are selectively expressed in the periosteum and contribute to fracture healing, while some markers, such as leptin receptor, do not appear until adulthood (4, 11-14). There is yet to be consensus about which markers to use and in what settings to identify mouse SSPCs. In humans, data is more limited as most studies select cells of interest based on adherence rather than using markers expressed in vivo that allow prospective isolation and localization. Chan and colleagues have characterized a fetal human skeletal stem cell (hSSC) population and various committed progenitor populations based on cell surface markers including CD73, PDPN, CD164 and CD146 (15). These markers and populations were initially identified and characterized in fetal growth plate tissue, then applied to various other tissues, including adult bone marrow and articular cartilage (15, 16). Other studies have used CD146, CD271, or PDGFRα and CD51 to isolate bone marrow populations with SSC properties (17-21).

The periosteum is essential for fracture repair (22, 23). Functionally, the periosteum niche differs from the bone marrow compartment in that it can form a callus containing both fibrocartilage and bone, but it does not provide a hematopoietic niche (12, 24). Many groups, including ours, have shown that the mouse periosteum contains high proportions of long-term skeletal stem cell populations with greater clonogenicity than the bone marrow (4, 12, 25, 26). Human periosteum has been utilized in various in vitro studies, but few have examined cell populations in freshly isolated periosteum. Debnath et al. briefly reported a periosteal stem cell population and two downstream progenitor populations identified in adult humans using analogous markers to what they identified in mice (10, 12).

To date, we have limited knowledge of SSPC populations in adult human tissue and how they compare between different skeletal compartments. In this study, we evaluated adult hSSPC populations across matched skeletal tissues, including periosteum, articular cartilage, and the bone marrow compartment, including central bone marrow and endosteum. We demonstrate vast differences in stromal populations in the different tissue compartments based on cell surface marker expression. In addition, we have evaluated the expansion and differentiation potential of selected periosteal populations in vitro.

## Results

### High-dimensional analysis reveals the heterogeneity of hSSPC populations in adult skeletal tissues

We performed spectral flow cytometry analysis of four skeletal tissues isolated with a panel containing 16 markers proposed to separate skeletal populations (Table 1). Periosteum, macroscopically normal articular cartilage, trabecular bone-associated cells (called endosteum) and bone marrow were isolated from the femoral head of 21 patients undergoing hip arthroplasty due to osteoarthritis (Figure 1A, Table 2). FCS files for the full dataset can be found at FlowRepository ID FR-FCM-Z5U6. Figure 1B illustrates the initial gating to generate skeletal lineage populations. We excluded hematopoietic, endothelial, erythroid cells and granulocytes using lineage (Lin) markers CD45, CD31, CD235a and CD15. We confirmed that all cells capable of forming fibroblastic colonies (CFU-F) resided within the Lin-fraction (Figure 1-figure supplement 1A). To look for high-level differences in the Lin-populations of the four tissues, we used the viSNE algorithm (27), followed by FlowSOM (28). We identified tissue-specific or selective clusters with hierarchical clustering (Figure 1C, Figure 1-figure supplement 2). CD200^hi^CD34^hi^ cells were predominately present in the periosteum (77.3% of the total CD200^hi^CD34^hi^), as were CD90^hi^ cells (54.8%). One cluster strongly overlapped with the hSSC population described by Chan (15), and 83.8% of this cluster was in the articular cartilage. The CD24+ clusters were specific to bone marrow (80%) and endosteum (17%), while the CD105^hi^CD164^hi^ cluster was almost exclusively present in the bone marrow (98.1%).

**Table 1.** Spectral flow cytometry panel.

| Antigen | Fluorophore | Clone | Cat # | Manufacturer | Dose<sup>1</sup> | Expression<sup>2</sup> | Reference<sup>3</sup> |
| --- | --- | --- | --- | --- | --- | --- | --- |
| ALP | BV650 | B4-78 | 742712 | BD Biosciences | 0.6 | Osteoblasts | (52) |
| CD105 | BV750 | 266 | 747107 | BD Biosciences | 0.6 | Endothelial cells, stromal cells | (12, 53) |
| CD106 (VCAM1) | BV421 | STA | 305815 | Biolegend | 0.6 | Inflamed endothelium, macrophages | (54) |
| CD133 | BB700 | W6B3C1 | 747638 | BD Biosciences | 0.6 | Cancer stem cells | (55) |
| CD141 | BB515 | 1A4 | 565084 | BD Biosciences | 1.25 | Dendritic cells, blood and lymphoid tissues, | (56, 57) |
| CD146 (MCAM) | BV605 | P1H12 | 361024 | Biolegend | 0.3 | MSCs, pericytes, endothelial cells | (1, 17, 19) |
| CD164 | PE | 67D2 | 130-126-518 | Miltenyi Biotec | 1.25 | HSC, SSC | (15, 58) |
| CD200 | APCFire750 | OX-104 | 329224 | Biolegend | 0.6 | BMSCs, periosteal progenitors | (12) |
| CD24 | BV711 | ML5 | 311135 | Biolegend | 0.6 | B cells, T cells, cancer stem cells | (59, 60) |
| CD26 (DDP4) | PE-Cy5 | BA5b | 302708 | Biolegend | 0.15 | Placental myofibroblasts | (37, 61) |
| CD271 (NGFR) | PE-CF594 | C40-1457 | 563452 | BD Biosciences | 0.6 | Neural crest-derived cells, MSCs | (17, 62) |
| CD34 | PerCP | 581 | 343520 | Biolegend | 2.5 | Various stem cells | (32, 35) |
| CD51 | APC | APC | 304415 | Biolegend | 0.6 | Broad stromal expression | (18) |
| CD73 | BV785 | AD2 | 344028 | Biolegend | 0.3 | MSCs, endothelial cells | (15, 53) |
| CD90 (Thy1) | Alexa Fluor 700 | 5E10 | 328120 | Biolegend | 0.6 | MSCs, HSCs, cancer stem cells, fibroblasts, pericytes | (53) |
| PDGFRα | PE-Cy7 | 16A1 | 323507 | Biolegend | 0.6 | Mesenchymal cells | (18) |
| PDPN | PerCP-eFluor 710 | NZ-1.3 | 46-9381 | eBioscience | 2.5 | Stromal cells, lymphatic endothelium | (15, 63) |
| **Lineage Markers** |  |  |  |  |  |  |  |
| CD235a | Pacific Blue | HI264 | 349107 | Biolegend | 0.3 | Erythroblasts/cytes |  |
| CD15 | Pacific Blue | W6D3 | 323022 | Biolegend | 0.6 | Granulocytes |  |
| CD45 | Krome Orange | J33 | B36294 | Beckman Coulter | 0.6 | Pan-haematopoietic marker |  |
| CD31 (PECAM1) | BV480 | WM59 | 566195 | BD Biosciences | 0.6 | Endothelial cells |  |

1. Antibody dose in $\mu\text{l}$ /tube. Staining carried out in 100 $\mu\text{l}$ volume 2. Selected cell types where expression is commonly reported 3. Previous use for separating skeletal or mesenchymal populations ALP, alkaline phosphatase; HSC, hematopoietic stem cells; MSC, mesenchymal stem cells; SSC, skeletal stem cells

**Table 2.** Patient demographics.

| Study | n | Sex | Age (Range) |
| --- | --- | --- | --- |
| Flow analysis | 21 | 11F, 10M | 65.3 (41-89) |
| Cell sorting | 11 | 6F, 5M | 68.7 (56-86) |
| Histology | 3 | 2F, 1M | 68.0 (53-77) |

**Figure 1.**
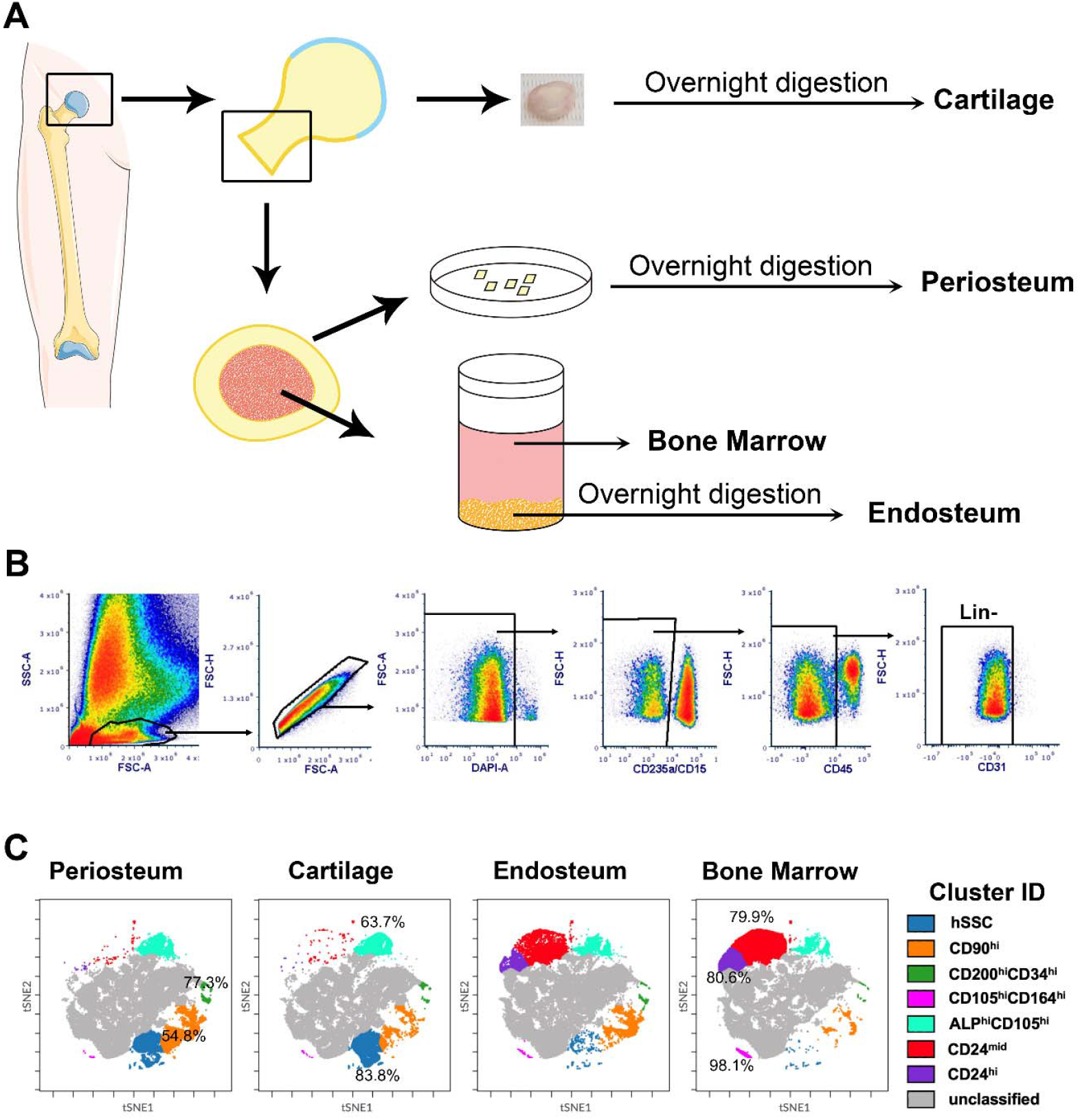
High level flow cytometry of skeletal tissues. (A) Skeletal tissue dissection and cell isolation protocol. (B) Gating strategy to identify lineage negative (Lin-) skeletal cells. (C) High-level analysis was performed using viSNE and FlowSOM. viSNE was run with equal sampling (325,000 events) of the concatenated Lin-populations for each tissue. FlowSOM clusters with clear differences between tissues, along with markers that define them are shown. The percentages indicate the proportion of events in that cluster within the indicated tissue.

We compared the expression of individual markers within Lin-populations in different tissues (Figure 2). None of the markers tested were specific to the Lin-population as they were detected in at least some of the Lin+ subsets (Figure 2-figure supplement 1). The periosteum showed the highest expression of CD90 (or THY1) and CD34, a stem cell marker in hematopoietic and endothelial cells (Figure 2A-C). Bone marrow showed the highest expression of perivascular marker CD146, consistent with its presence on sinusoidal walls, although it remained rare in all tissues (Figure 2D) (19). CD24, used to separate marrow stromal populations in mice (29), was highly expressed in bone marrow and very rare in periosteum (Figure 2D-E). In the endosteum, CD24+ cells had limited ability to form CFU-F (Figure 1-figure supplement 1B), suggesting they primarily represent a mature marrow stromal population rather than progenitor cells. Articular cartilage showed significant enrichment of many of the markers evaluated, including CD164, PDGFRα, PDPN, and ALP, a mature osteoblast marker (Figure 2F-G). Patient gender did not significantly affect the frequency of most markers analyzed with the exception of CD90 in periosteum which showed higher frequency in males (p=0.0074, see Figure 2 – Source data 1). Expression of some markers appeared to be affected by patient age, although only in selected tissues. These included CD26 in bone marrow which tended to increase with age, age-related increases in ALP in endosteum and periosteum, and CD24 in endosteum which decreased with age (Figure 2-figure supplement 2).

**Figure 2.**
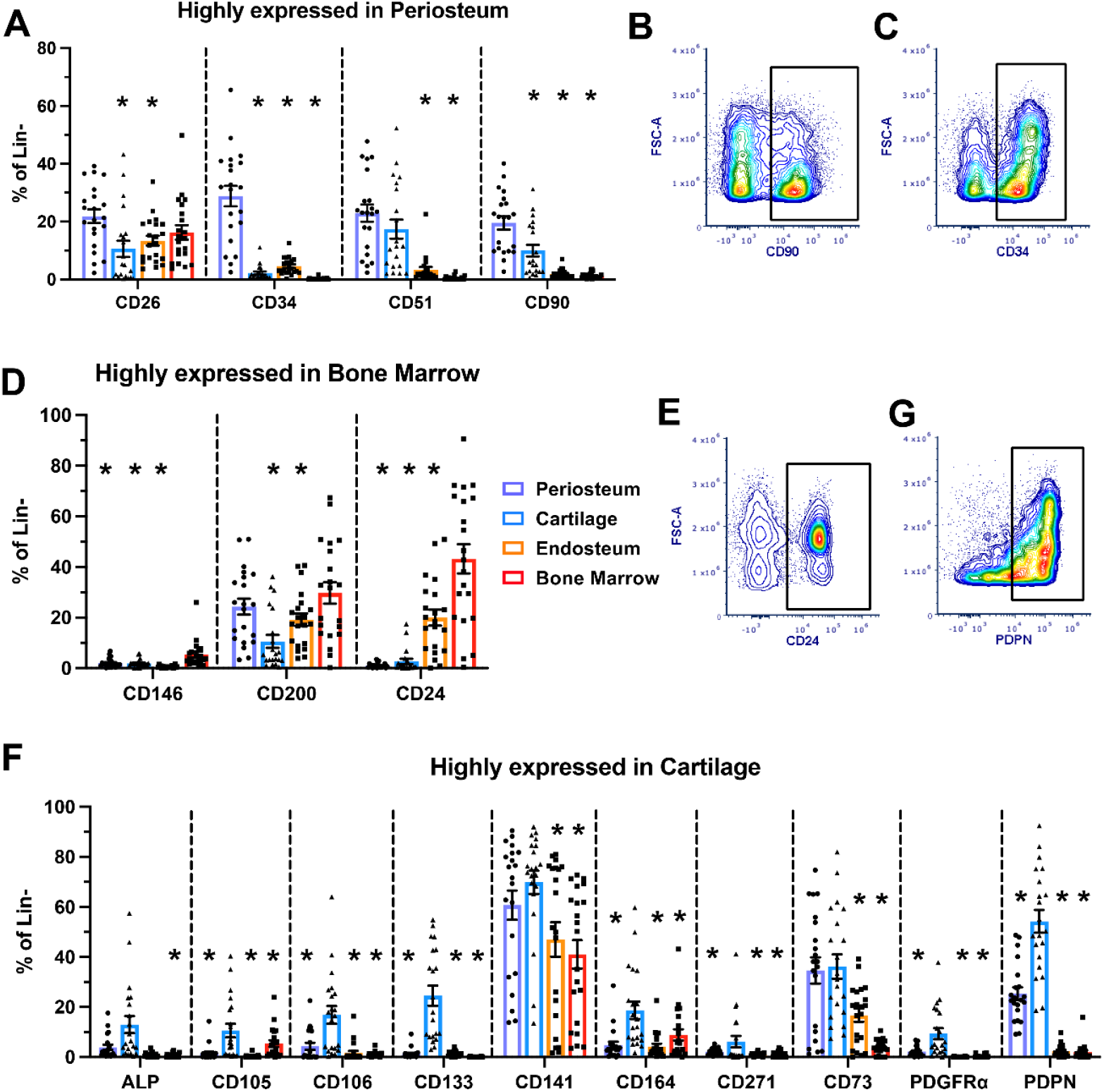
Marker expression varies in different skeletal compartments. Expression of individual markers in the lineage negative (Lin-) populations isolated from different tissues using a multi-colour flow panel. (A) Markers that are highly expressed in the periosteum, see D for legend for all graphs. Representative density plot of CD90 (B) and CD34 (C) in the periosteum. (D) Markers that are highly expressed in the bone marrow. Representative density plot of CD24 (E) in the bone marrow. (F) Markers that are highly expressed in the cartilage. Representative density plot of PDN (G) in the cartilage. n=21. *p<0.05 compared to the tissue indicated in the graph title, one-way ANOVA with Dunnett’s post hoc test.

We evaluated previously reported human skeletal stem cell stains in our dataset. The hSSC, as defined by Chan et al. (15) (Lin-CD146-PDPN+CD164+CD73+), but not their proposed downstream progenitors, appear as a cluster on our viSNE analysis (Figure 3A). hSSCs showed clear separation making gating straightforward in most cartilage samples but not in other tissues (Figure 3B-C). The proportion of hSSC was three times higher in the cartilage (7.8% of Lin-) compared with the periosteum (2.9%, Figure 3D). hSSCs were very rare, or in some patients absent, in the endosteum (0.13%) and bone marrow (0.03%). Osteoprogenitors (hOP, Lin-PDPN-CD146+) were most common in the bone marrow (5.4%), and rare in other tissues (Figure 3E), which does not align with the known functionality of these tissues and the extensive data indicating that osteoprogenitors reside near the bone surface (21, 30, 31). Bone, cartilage and stromal progenitors (hBCSP, Lin-CD146+PDPN+) were rare in all tissues (<3%), while chondroprogenitors (hCP) showed the highest frequency in cartilage (13.7%), followed by the periosteum (7.6%, Figure 3F-G). None of the periosteal stem and progenitor populations hPSC (Lin-CD90-CD105-CD200+), hPP1 (Lin-CD90-CD105-CD200-), and hPP2 (Lin-CD90-CD105+) proposed by

**Figure 3.**
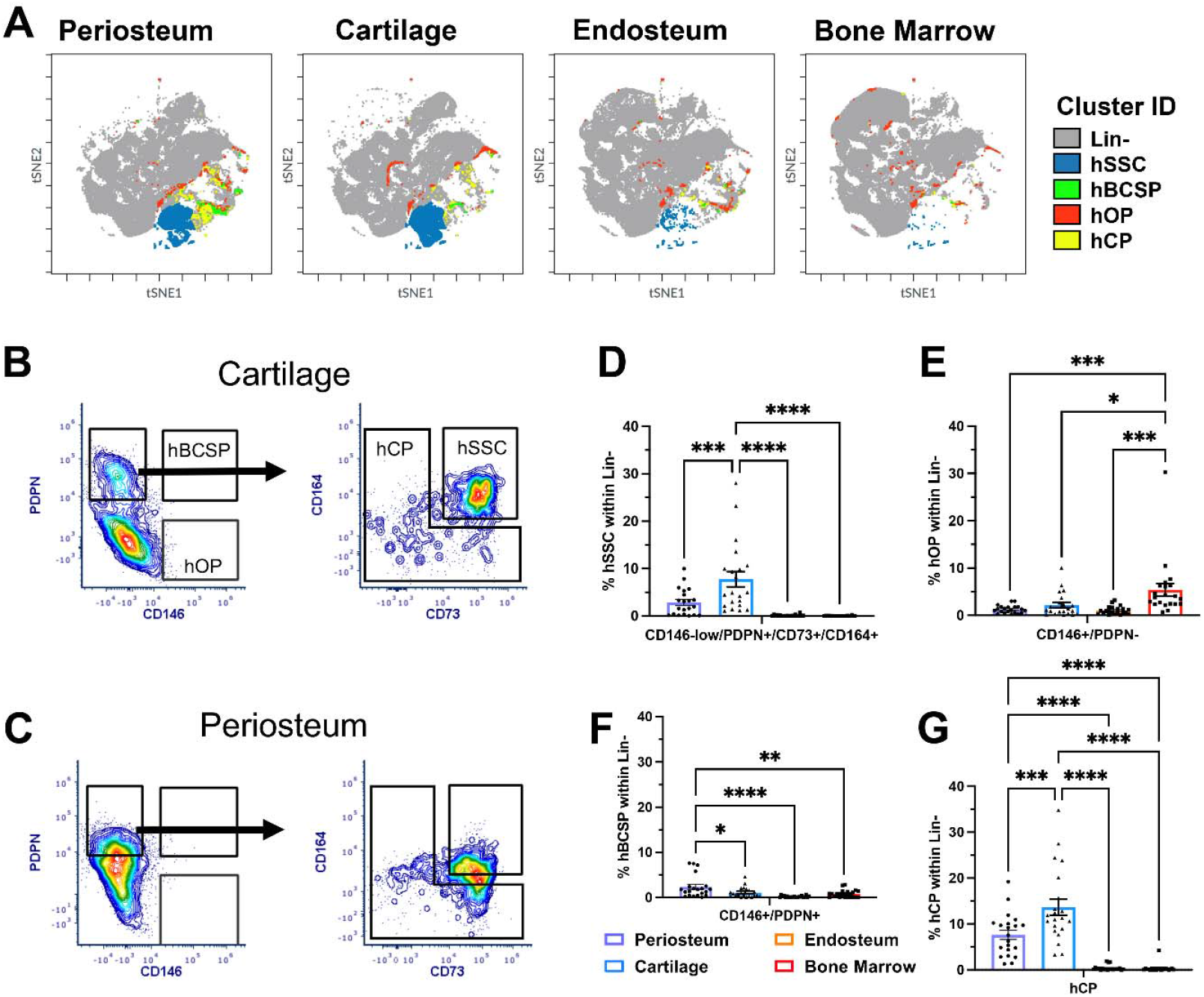
Distribution of hSSC and downstream populations in adult tissues. (A) Skeletal stem cell (hSSC), bone cartilage stromal progenitor (hBCSP), osteoprogenitor (hOP), and chondroprogenitor (hCP) populations described in (15) overlaid onto viSNE plots of the four tissues analyzed. Gating to identify these populations in (B) cartilage, and (C) periosteum. Frequency of (D) hSSC, (E) hOP, (F) hBSCP, and (G) hCP in Lin-fractions of periosteum, cartilage, endosteum, and bone marrow samples, n=21. *p<0.05, **p<0.01, ***p<0.001, ****p<0.0001, one-way ANOVA with Turkey’s post hoc test.

Debnath et al. (12) presented as a cluster on our viSNE plots except for in the bone marrow where hPSC overlapped with the CD24+ cluster (Figure 3-figure supplement 1A). The hPSC population was present in the periosteum, but rare in other tissues, but hPP1 was very common in all tissues (>65% of Lin-), suggesting this set of markers may not be suitable for non-periosteal tissues (Figure 3-figure supplement 1).

### Enrichment of SSPCs using CD90

We evaluated the growth and differentiation potential of various prospectively isolated cell populations using in vitro assays (Figure 4A). Initially, we evaluated CFU-F formation in each tissue’s total Lin-populations. 1.7±0.16% of Lin-periosteum cells formed CFU-F, around 5-fold higher than the CFU-F in endosteal preparations (0.34±0.23%, Figure 4B). Surprisingly, CFU-F frequency in articular cartilage, 2.3±0.24%, tended to be higher than periosteum despite the inability of cartilage to heal; however, cartilage CFU-F were consistently smaller and contained fewer cells than matched periosteum samples. Bone marrow did not form CFU-F despite being plated at 10x higher density, consistent with previous reports from humans and mice indicating that CFU-Fs and SSPCs primarily reside near the bone surface (4, 15, 21, 31). Next, we attempted to enrich for periosteal CFU-F using markers highly expressed in the periosteum with clear separation allowing straightforward and reproducible gating. We focused on CD90 as this marker enriched for CFU-F capable of multi-lineage differentiation in our mouse studies (4). A subset of CD90+ cells in endosteum and bone marrow were capable of CFU-F formation, while CD90-cells were not (Figure 4-figure supplement 1A-D). In cartilage, CD90 did not enrich for CFU-F. Expanded CD90+ colonies showed varied in vitro differentiation potential (Figure 4-figure supplement 1E-F). In the periosteum, we split the Lin-population by combining CD90 with CD73, CD34 and CD26 (Figure 4C-E). There is partial but not complete overlap in the expression of these markers (Figure 4-figure supplement 1G). The CD90+CD34+ population was the only one showing significant CFU-F enrichment compared to total Lin-. We then further characterised individual clones from populations with consistent CFU-F formation. Most colonies selected for passaging were capable of further expansion, but CD90+CD34+ cells expanded more rapidly than the other populations (Figure 4F-G). Most expanded colonies were also capable of differentiation, but notably, 100% of the 11 CD90+CD34+ clones, as well as all 12 of CD90+CD26+ clones (each derived from 3 patients) were capable of differentiating into osteoblasts, adipocytes and chondrocytes under permissive conditions. CD90-CD73+ cells from 2/3 patients underwent spontaneous adipogenesis prior to the addition of adipogenic medium while retaining osteogenic and chondrogenic potential in most cases.

**Figure 4.**
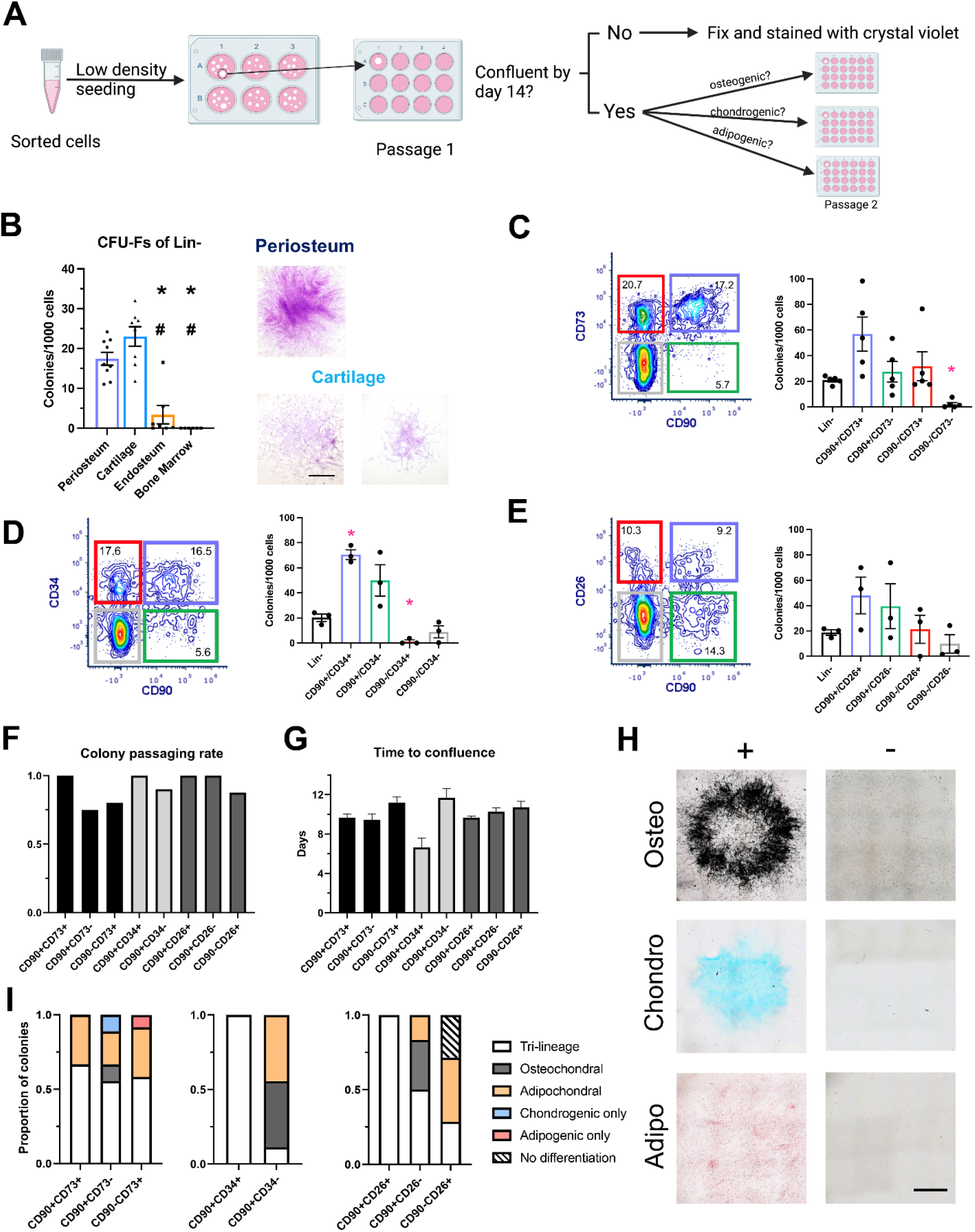
In vitro characterization of potential skeletal stem cell populations. (A) Experimental design for single colony functional analysis experiments (created with BioRender). (B) Colony-forming unit fibroblast (CFU-F) frequency in lineage negative (Lin-) skeletal cells from different tissues, n=6-9. Representative colonies stained with crystal violet are shown, scale bar = 500μm. CFU-F frequency in periosteal (C) CD90/CD73 subpopulations, n=5, (D) CD90/CD34 subpopulations, n=3, and (E) CD90/CD26 subpopulations, n=3. (F) The proportion of single colonies from each population reaching confluence within 14 days, 3-4 colonies were picked from each population per sample, n=2-4 donors/population. (G) Time for the picked colonies to reach confluence. (H) Examples of positive and negative staining for von Kossa (osteo), alcian blue (chondro), and oil red O (adipo) in individual expanded colonies, scale bar = 0.25cm. (I) Differentiation potential of expanded colonies when grown under permissive conditions. (B) *p<0.05 compared to periosteum, #p<0.05 compared to cartilage, one-way ANOVA with Turkey’s post hoc test. (C-E) *p<0.05 compared to Lin-, one-way repeated measures ANOVA with Dunnett’s post hoc test.

### Localization of CD90+CD34+ cells in the periosteum

We localized the CD90+CD34+ populations in situ with immunostaining on femoral neck sections. Consistent with the flow cytometry data, CD90+ cells were most abundant in the periosteum. CD90+CD34+ cells were mainly located in the periosteum, but comprised at least two populations (Figure 5A-C, H). Perivascular (lectin-adjacent) cells with robust CD90 staining were present in the outer layer of the periosteum (Figure 5B). A second non-perivascular CD90+CD34+ population was evident in the inner cambium region of the periosteum where periosteal stem cells are proposed to reside (Figure 5C). CD73+ immunostaining was rare in the periosteum in contrast to our flow data, and CD73+CD90+ cells were most abundant near the endosteal bone surface (Figure 5D-G, I). The CD90+CD34+ marker combination, therefore, has utility using both flow cytometry and histology; however, additional markers are needed to refine these cells to a more homogeneous population.

**Figure 5.**
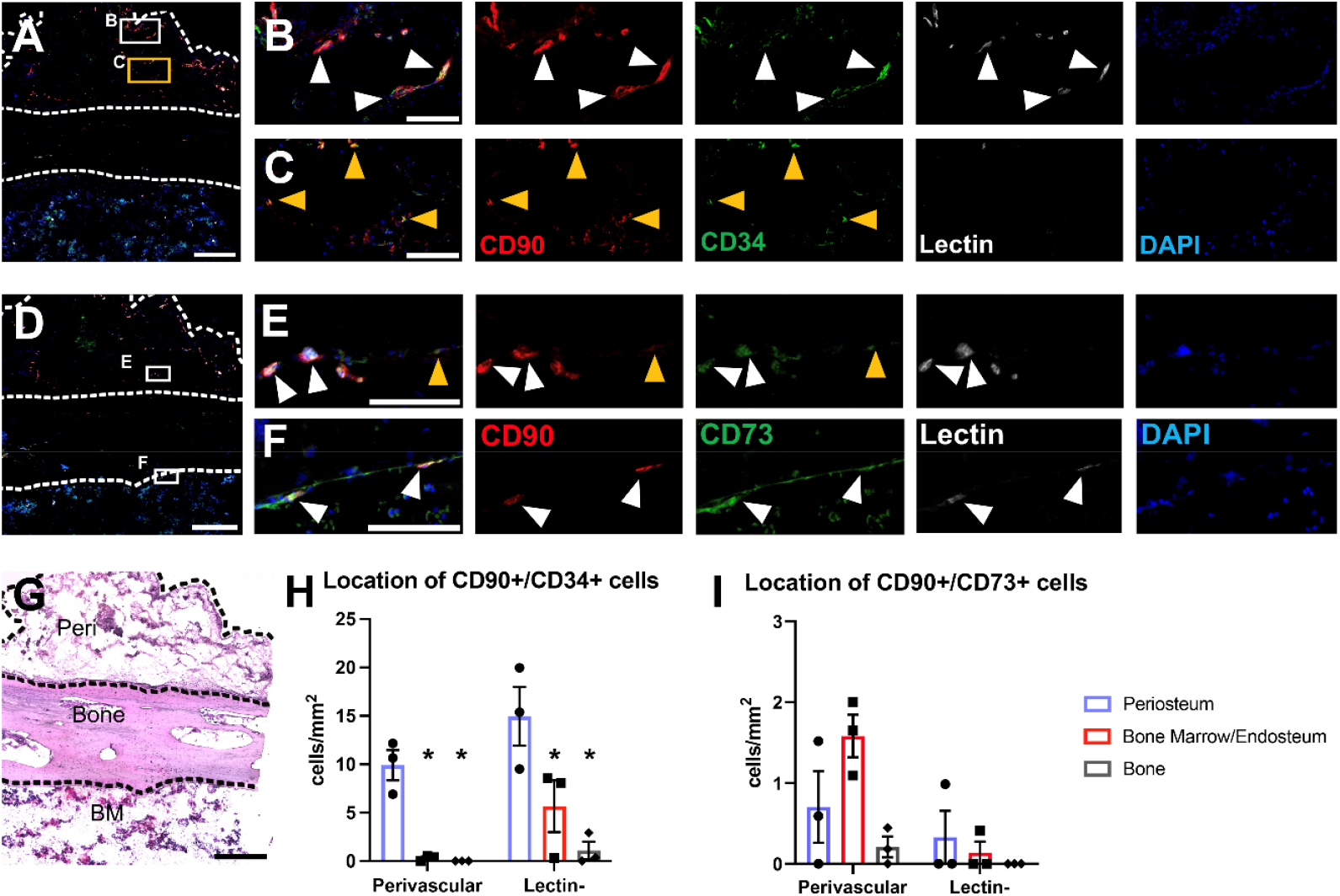
Two subsets of CD90+CD34+ are located in the periosteum. Immunostaining was performed on femoral neck sections of 3 individuals. (A) Localization of CD90+CD34+ subsets including (B) perivascular, and (C) non-perivascular periosteal populations. (D) CD90+CD73+ subsets in (E) the inner periosteum, and (F) the endosteal surface. Nuclei are counterstained with DAPI (blue), and endothelial cells are stained with lectin (white). White arrowheads indicate dual stained cells adjacent to vasculature, yellow arrowheads indicate non-perivascular dual positive cells. (G) Hematoxylin and eosin staining of an adjacent section. BM, bone marrow; peri, periosteum. Quantification of (H) CD90+CD34+, and (I) CD90+CD73+ cells in different regions of the bone. Scale bar = 500μm (A, D, G) or 100μm (B-C, E-F). *p<0.05 compared to periosteum, 2-way repeated measures ANOVA with Dunnett’s post hoc test.

## Discussion

Data generated with single-cell techniques indicate that there is a great deal of cellular diversity in skeletal tissues. Using spectral cytometry of cell surface markers, we have demonstrated that the Lin-skeletal populations resident in the periosteum, the bone marrow compartment and articular cartilage are very different with some cell populations restricted to certain tissue types. Presumably, these differences would be amplified further with higher-resolution techniques like single-cell RNAseq. While the idea of universal markers or marker combinations is appealing, abundant evidence shows that populations expressing similar markers sourced from different tissues have very different characteristics. For example, CD146+ cells from various skeletal and non-skeletal tissues form CFU-F, but only bone marrow-derived cells form ossicles containing marrow, while periosteal cells make ossicles without marrow infiltration (5). Similarly, markers including CD271 and PDGFRα that enrich for bone marrow CFU-F show very different abundance in fetal compared to adult bone marrow, and the combinations that show the best enrichment of stem cell populations appear to vary in different developmental stages (18, 20). Despite the clear differences between tissues, we saw limited influence of gender and age on individual marker expression in our dataset. Notably, all patients in our study were over 40, and almost all women would have been postmenopausal, so larger cohorts including younger patients would be required to thoroughly address the effect of age on resident skeletal lineage populations.

Many studies on SSPCs use plastic adherence, or adherence to fibronectin-coated plates for chondroprogenitors, but numerous studies have shown that the cell surface phenotype of various mesenchymal progenitor populations is altered by attachment, in vitro culture conditions, and passaging (32-37). Therefore, it is important to identify functionally different SSPCs and define their origins in vivo. Consistent with recent studies in mice, our results demonstrate that hSSPCs are enriched in the adult periosteum compared to the bone marrow and endosteal compartments (4, 12, 14, 25, 26). In line with other studies, we found that CFU-Fs were very rare in total Lin-bone marrow, and sorting on the basis of rare markers, CD90-based enrichment in this instance, was necessary to detect any colony formation (15, 21). In the periosteum, most cells capable of CFU-F formation were CD90+, and the CD90+CD34+ fraction was enriched 3.5-fold for CFU-F. These colonies also showed consistent expansion and multi-lineage differentiation on a clonal basis, suggesting that periosteal stem cells reside within the CD90+CD34+ fraction. This contrasts with the results reported by Debnath et al., where only CD90-cells were considered to be periosteal stem and progenitors (12). Notably, there were CD90-cells capable of CFU-F formation, particularly CD90-CD73+ cells. Some clones from this population also demonstrated tri-lineage differentiation, but notably, many formed adipocytes spontaneously, which is surprising given that in vivo the periosteum does not contain adipocytes. We have noted that mouse periosteal cultures also form adipocytes readily, often more rapidly than bone marrow stromal cells, suggesting that the in vivo periosteal environment may actively inhibit adipogenic differentiation (13, 38). Overall, the vast majority of colonies chosen for clonal analysis in this study were capable of further expansion and some form of differentiation, which contrasts with our mouse studies where periosteal CFU-F usually have limited ability to expand in vitro without the addition of growth factors, in agreement with other studies (15, 39). In mice, prospectively isolated CD90+ cells are described as committed osteoprogenitors rather than stem cells, but most of this data was generated using cells from embryonic or early postnatal donors, suggesting the functionality of CD90+ cells may be different in development compared to adulthood (10, 40). Notably, CD90+CD34+ cells also comprise a clear population in synovial fibroblasts and show multi-lineage differentiation potential, at least in bulk cultures (41, 42). While CD90+CD34+ cells are effectively enriched for cells with stem cell properties, at least in vitro, additional markers are still required to separate the population further, particularly the perivascular population from the non-perivascular subset resident in the cambium layer of the periosteum. The cambium layer is generally proposed to contain the stem cell niche, and cells that contribute to fracture callus tissue formation are usually close to the bone surface in resting periosteum, although it is difficult to identify the layers in many parts of the mouse skeleton (12, 14, 43, 44). Further refinement of this population using additional markers will be addressed in future studies.

Articular cartilage showed high expression of many putative SSPC markers tested, including high expression of ALP chosen as a mature osteoblast marker. Previous studies have also reported strong expression of selected markers, including CD73 and CD106 in articular cartilage, and two studies show around 25% CD90+ cells, which is higher than in our study (10% of Lin-) (34, 45, 46). Collectively, these data suggest that some progenitor markers are expressed in mature chondrocytes, although in some cases, also chondroprogenitors (34). Our data suggest that CD73 and PDPN are expressed in a large portion of chondrocytes. Surprisingly, cartilage showed high overall CFU-F formation with similar or even higher levels than in periosteum. The size and morphology of colonies forming from articular cartilage suggested a slower growth rate and potentially more limited expansion potential than periosteal cells, although we have not examined this question systematically. Cartilage contained the highest proportion of hSSCs, as defined by Chan et al., and bright staining for several markers used to define hSSCs (15). Given that this stain was developed using growth plate tissue, it is perhaps not surprising that it appears most suited to cartilage. Our data suggest that further validation of the hSSC hierarchy is required in adult tissues.

The major limitation of this study is the use of tissue from femoral heads of osteoarthritis patients. This limited the source of periosteum to the femoral neck, which has partial periosteal coverage and does not produce fracture callus well (47). However, it did enable us to analyze a large cohort that included both genders and source our cells from a consistent location. Future studies will be required to confirm replication with different anatomical sources of periosteum. The cartilage used for these analyses was macroscopically normal. While this means it experienced an osteoarthritic environment, there are clear differences in various cellular phenotypes between chondrocytes obtained from macroscopically normal and matched damaged cartilage, indicating they retain some features of healthy cartilage (48, 49). Some potential SSPC markers retain similar distribution and frequency in osteoarthritis, while others change in disease (34). Tissues adjacent to osteoarthritic joints, including the bone, are affected by osteoarthritis, which could potentially affect the SSPC and stromal population frequencies, although the site of marrow collection was some distance from the subchondral region that has overt changes. Finally, we restricted our analysis of expansion and differentiation potential to in vitro assays, however we performed these on a clonal basis which meant we observed a variety of differentiation potentials.

In conclusion, we have demonstrated that adult human periosteum is enriched for SSPCs compared to the bone marrow compartment, but perhaps not articular cartilage despite its vastly superior regenerative capacity. Our data suggest that these different tissue compartments have substantial differences in the skeletal/stromal cell makeup and that different combinations of markers will be required to identify SSPCs in periosteum, the bone marrow compartment, and articular cartilage. Our results do not exclude the possibility of multiple separate stem cell pools with different functionality, although it will be challenging to interrogate in vivo function in humans. Finally, we demonstrated that a subset of CD90+CD34+ cells in the periosteum have characteristics of skeletal stem cells.

## Methods

### Collection of adult human skeletal samples

Collection and use of human tissue were approved by The New Zealand Northern A Health and Disability Ethics Committee (NTX/05/06/058/AM15) and all participants provided written informed consent. Femoral heads were collected from patients undergoing hip arthroplasty for osteoarthritis at Auckland City Hospital, Auckland, New Zealand (Table 2). Specimens were kept in sterile saline at 4°C for no longer than 6 h before dissection.

### Histology

The cortical ring was sawed from the femoral neck and trimmed to approximately 1cm^2^ before fixing with 4% paraformaldehyde (Sigma, NZ) at 4°C for 5-7 days. Samples were then decalcified with 14% ethylenediaminetetraacetic acid (EDTA) changed weekly for six months, dehydrated in 30% sucrose overnight, and stored frozen until sectioning. The tissue was embedded in cryomatrix (ThermoFisher) and 7μm cryosections were obtained on a cryostat (CryoStar, Leica Microsystems, Wetzlar, Germany) with a tape transfer system (Section-lab, Hiroshima, Japan) as previously described (50). For immunofluorescent staining, sections were permeabilized with 0.3% Triton X in PBS for 15min, followed by 1h blocking in 5% bovine serum albumin (BSA) with 10% normal goat serum (NGS) in 0.1% Tween 20/PBS (PBST) at RT. After blocking, sections were incubated with a primary antibody cocktail (Table 3) made up in 1% BSA/2%NGS/PBST (antibody diluent) at 4°C overnight. Where required, slides were washed and secondaries were incubated for 1h at RT. All sections were counterstained with DAPI before mounting with ProLong Diamond Antifade Reagent (ThermoFisher). For haematoxylin and eosin staining followed the manufacturer’s protocol (Section-lab). Slides were scanned using a Metafer4 Slide Scanning Platform (MetaSystems, Altlussheim, Germany) at 10X. Image analysis was performed using image J (Fiji). Regions of interest (ROIs) including the periosteal, bone, and endosteal/bone marrow regions were drawn according to the adjacent brightfield images. Standardized thresholds were manually set for each channel, and the fluorescent signal for each channel was measured in DAPI-positive nuclear regions after nuclei were separated with the watershed algorithm. We calculated cells that were positive for CD90, CD34, CD73, or lectin; double positive for CD90/CD34 or CD90/CD73; and triple positive for CD90/CD34/lectin or CD90/D73/lectin in different ROIs. One section/bone was analyzed.

**Table 3.** Reagents used for immunostaining.

| Antigen | Conjugate | Clone | Cat# | Manufacturer | Dilution |
| --- | --- | --- | --- | --- | --- |
| CD90 | APC-Cy7 | 5E10 | 328131 | Biolegend | 1:50 |
| CD34 | PE/Dazzle 594 | 581 | 343534 | Biolegend | 1:25 |
| CD73 | - | AD2 | 344002 | Biolegend | 1:100 |
| UEA-1 Lectin | Biotin |  | L8262 | Sigma | 1:500 |
| Goat anti-mouse Alexa Fluor 594 |  |  | A-11032 | ThermoFisher | 1:500 |
| Streptavidin Alexa Fluor 488 |  |  | S11223 | ThermoFisher | 1:500 |

### Cell isolation from skeletal tissues

Periosteum, cartilage, endosteum, and bone marrow were isolated from the femoral head of the same patient (Figure 1A). The periosteum was scraped off the cortical ring and minced. Macroscopically undamaged articular cartilage was dissected and cut into approximately 2mm^2^ pieces. Trabecular bone was extracted from the femoral head and washed with cold PBS, the washes were collected as bone marrow, and the cleaned trabecular bone was further washed and minced. All tissues except bone marrow were incubated with 5mL/g tissue of 1mg/mL collagenase P (Cat: 11-213873001, Sigma-Aldrich) in αMEM 10% fetal bovine serum (FBS), at 37°C, 100rpm overnight (<15h). Following digestion, cells were filtered through a 70μm cell strainer (Falcon), washed with PBS, and the pellet was resuspended and incubated with red blood cell lysis buffer (155mM NH_4_Cl, 10mM KHCO_3_, 0.1mM EDTA in H_2_O) for 1min then washed and resuspended in staining medium (SM, 2% FBS, 1mM EDTA in PBS).

### Flow cytometry and cell sorting

We used Panel One reported in Boss et al. (37) and similar panels used for the analysis of adipose tissue as the backbone of our stain. The panel is shown in Table 1. For spectral flow cytometry, cells were blocked with human TruStain FcR Blocking Reagent (Biolegend, USA) and True-Stain Monocyte Blocker (Biolegend, USA) before staining with antibody cocktails with Brilliant Stain Buffer (BD Biosciences, US) in SM. For cell sorting, cells were stained using antibody cocktails in SM. Dead cells were excluded using DAPI (50ng/ml final concentration).

Spectral flow cytometry analysis was performed on a Cytek Northern Lights instrument with three lasers. Each experiment included an unstained control using unlabeled freshly isolated periosteal cells for background observation and establishing gates. Spectral unmixing was calculated with a standard reference control library and freshly prepared unstained controls by SpectroFlo Software Package (Cytek Biosciences, USA). To build the reference library, periosteal cells were stained with each antibody used in the panel, but where the interested populations were dim or rare on the periosteum, other skeletal cells or beads (Compensation Plus (7.5μm) Particles Sets, BD Biosciences) were used instead.

Cell sorting was performed on a BD FACS Aria II using panels that included antibodies listed in Table 4. Cells were collected into 1.5mL sterile screw-cap tubes containing 500μL αMEM 20% FBS.

**Table 4.** Additional antibodies for cell sorting.

| Antigen | Conjugate | Clone | Cat # | Manufacturer | Dose (µL/100µL) |
| --- | --- | --- | --- | --- | --- |
| CD235a | FITC | HI264 | 349104 | Biolegend | 2.5 |
| CD31 | FITC | WM59 | 303104 | Biolegend | 2.5 |
| CD45 | FITC | HI30 | 555482 | BD Biosciences | 10 |
| CD26 | PE | BA5b | 302706 | Biolegend | 0.15 |
| CD73 | PE-Cy7 | AD2 | 561258 | BD Biosciences | 2.5 |
| CD90 | APC-Cy7 | 5E10 | 328131 | Biolegend | 1 |
| CD90 | PE | 5E10 | 561970 | BD Biosciences | 0.25 |

### Data analysis for spectral flow cytometry

After unmixing, FCS files were exported and analyzed with FCS Express 7 and FlowJo v10.6.2 (BD Biosciences). Debris, doublets, dead cells, and Lin+ cells were excluded with serial gating (Figure 1B). High-dimensional analysis was performed on gated Lin-populations in Cytobank (Beckman Coulter). The analyses were applied to the four equal sampled (325,000 events) concatenated Lin-populations. We used the advanced t-Distributed Stochastic Neighbor Embedding (tSNE) algorithm, viSNE (3000 iterations) (27), followed by FlowSOM hierarchical clustering (28) with 15 clusters by the consensus metaclustering method.

### In vitro cell culture

Sorted cells were resuspended in αMEM 20% FBS and seeded at 20-50 cells/cm^2^ for periosteum and cartilage and 200-2000 cells/cm^2^ for bone marrow and endosteum for CFU-F assays. Lin+ cells were seeded at 2000-6000 cells/cm^2^. Cells were cultured in a 37°C humidified incubator with 5-6% oxygen and 5% CO_2_. Half and full medium changes were performed on day 4 and 7, respectively, and colonies were either counted manually under a microscope at 4x or terminated for staining on days 9-10. For clonal analysis of CFU-F, medium-large colonies with suitable separation and positioning were chosen for passaging. Cloning rings were positioned, and cells detached with accutase (Cat: A1110501-01, Gibco, ThermoFisher Scientific). Each colony was transferred to one well of a 12-well plate and cultured in αMEM 10% FBS until 80-90% confluence. Medium was changed twice weekly. Wells were terminated if they did not reach confluence by day 14. Confluent wells were detached with accutase, resuspended in αMEM 10% FBS, and then split into 3 wells for differentiation in 24-well plates. 70% of the cells were seeded for chondrogenesis as a 25μl spot; 15% for osteogenesis as a 20μl spot; and 15% for adipogenesis in 500μl. After 2h, wells with spots were topped up with 500μL αMEM 10% FBS. For chondrogenic differentiation, the day after seeding medium was changed into serum-free DMEM high glucose containing 50μg/ml ascorbic acid, 100nM dexamethasone, 1X sodium pyruvate, 1X ITS+1, 40μg/mL L-proline, and 10ng/ml TGF-β3, and the plates were incubated at 5-6% oxygen for 9 days (13). For osteogenic differentiation, the medium was changed to αMEM 5% FBS, 50μg/mL ascorbic acid-2-phosphate, 5mM β-glycerophosphate, 10nM dexamethasone the day after seeding, then cultured in normoxic conditions for 21 days. For adipogenesis, cells were cultured until confluence (3-5 days) then changed to DMEM/F12 10% FBS, 10μM insulin, 200μM indomethacin, and 1μM dexamethasone for 21 days (51).

Cells were washed with PBS and fixed in 10% formalin for 5 min prior to any stain. CFU-F plates that did not undergo differentiation, or expanded clones that did not reach confluence were stained with 0.05% crystal violet. For osteogenesis, cells were incubated with 1.25% silver nitrate for 30 min. For adipogenesis assessment, cells were washed with 60% isopropanol then stained with 0.21% Oil Red O in 60% isopropanol for 30 min. Chondrogenesis was observed with alcian blue staining. Fixed cells were washed twice with 3% acetic acid (pH 1.0) and stained overnight with 1% alcian blue 8GX (in 3% acetic acid, pH 1.0). All the excess stains were aspirated after staining. For von Kossa and ORO staining, cells were washed three times with water and air dried. For alcian blue staining, cells were rinsed briefly with 3% acetic acid (pH 1.0) followed by 3% acetic acid (pH 2.5), and air dried. Plates were imaged using a Nikon TE2000E inverted fluorescence microscope (Nikon, Japan) at 4x.

### Statistics

For the spectral flow cytometry study, we empirically selected a sample size of 20 with even sex distribution as data suitable for power calculations was not available. The actual number was increased to 21 due to sample availability. For the detailed analysis within the smaller or rare populations in the flow data, samples with less than 100 events were excluded. Statistical analysis was performed in GraphPad Prism. For the CFU-F data where matched samples were used, paired tests were performed.

Details of statistical analysis, including exact n values, are listed in figures or figure legends. Each graph is presented as the mean ± standard error of the mean (SEM) unless otherwise stated. p<0.05 was considered statistically significant unless otherwise stated.

## Acknowledgements

We thank the donors for donating their hips, Hula Polima, the surgeons, and the theatre staff for collecting tissues and coordinating their transfer to the university. We thank Marcus Ground for assisting with generating selected figures using Biorender. We thank Thaize Chrometon for assisting with flow cytometry data collection. This work has been supported by the Health Research Council of New Zealand Sir Charles Hercus Fellowship and the Maurice and Phyllis Paykel Trust project grant to BGM. YC is supported by a University of Auckland Doctoral Scholarship.

---

**Figure 1-figure supplement 1.**
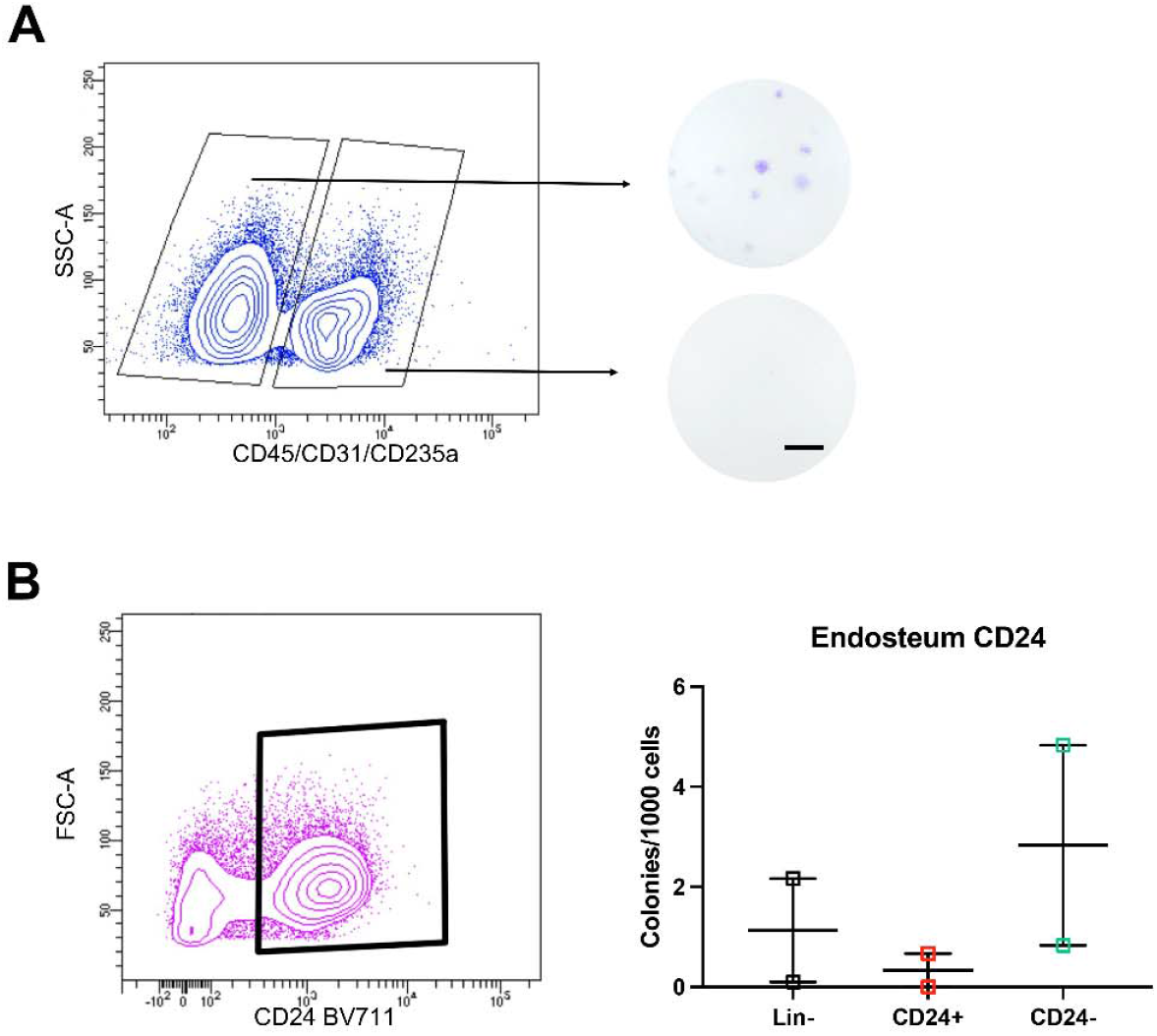
Colony forming potential of selected populations. (A) Density plot showing sorted periosteal lineage -/+ populations and their colony-forming ability confirmed with crystal violet. Representative of n=9. Scale bar = 0.5cm. (B) Density plot showing sorted CD24+ population and colony-forming unit fibroblast (CFU-F) frequency in endosteal cells, n=2.

**Figure 1-figure supplement 2.**
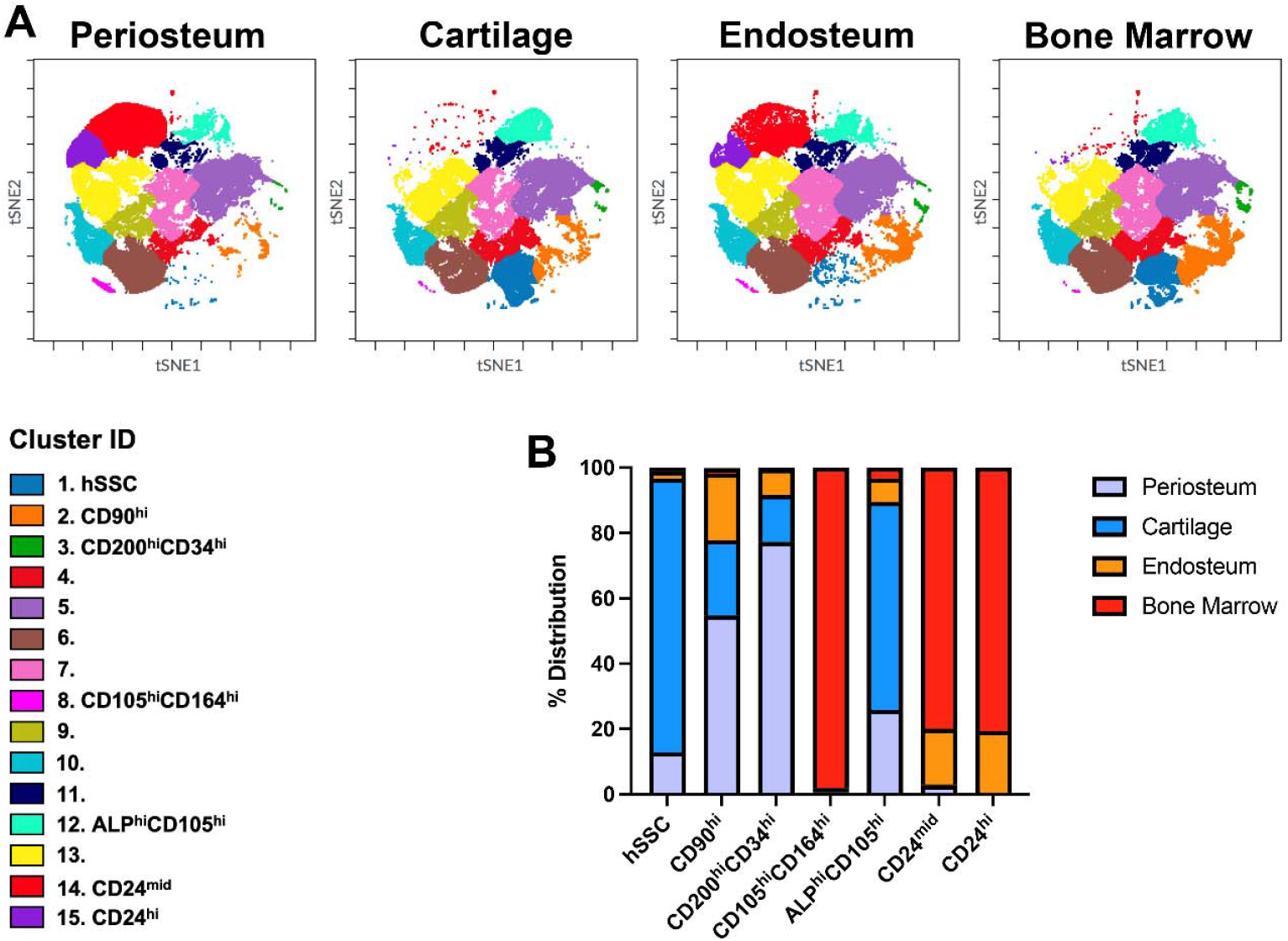
Full FlowSOM 15-cluster analysis of skeletal cells. (A) FlowSOM-defined clusters overlaid onto viSNE plots. Clusters with clear markers identified in Figure 1C are labelled, as are unclassified clusters. (B) Distribution of identified clusters in periosteum, cartilage, endosteum, and bone marrow. Note that equal numbers of events were initially included for each tissue.

**Figure 2-figure supplement 1.**
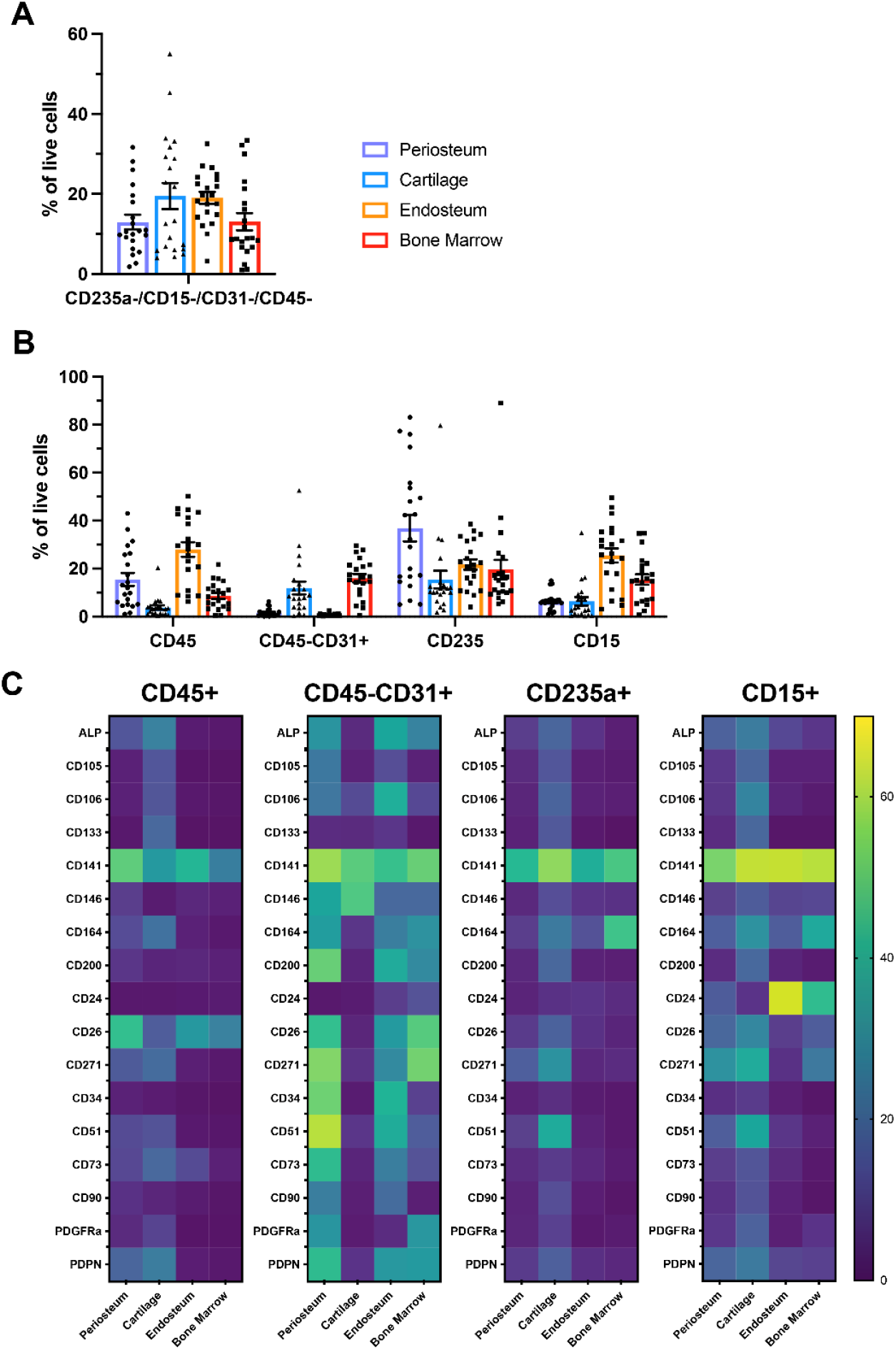
Expression of proposed skeletal stem and progenitor markers in non-skeletal lineages. (A) Proportion of lineage positive (Lin+) populations in samples analyzed using the 21-color panel. (B) Proportion of CD45+ (hematopoietic), CD45-CD31+ (endothelial), CD235a+ (erythroid), and CD15+ (granulocyte) populations of each tissue. (C) Heatmap of markers expressed on the different Lin+ populations of each tissue indicating average % positive cells for each marker across the 21 patient samples, data are shown as means.

**Figure 3-figure supplement 1.**
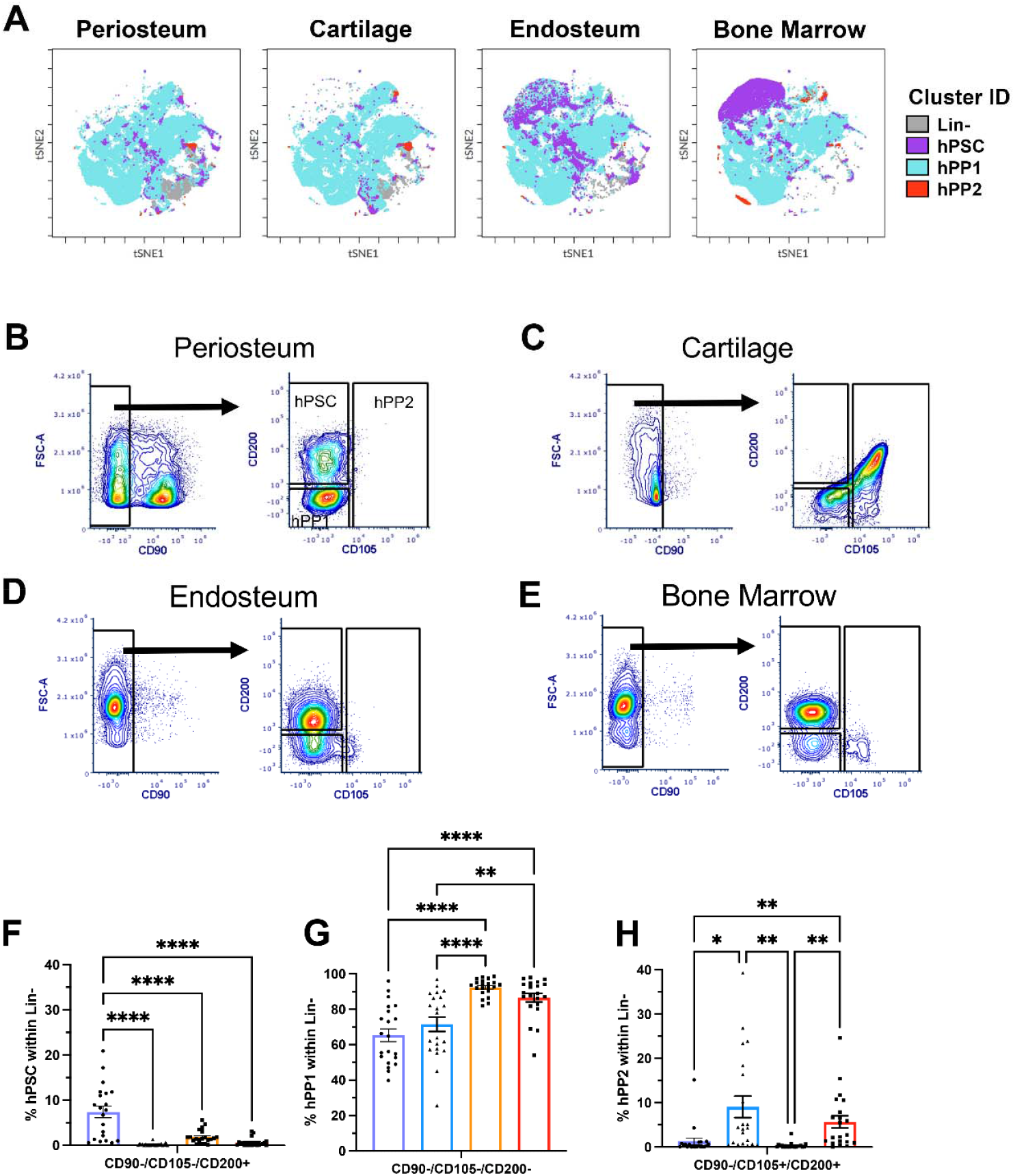
Human periosteal stem cell stain in different adult tissues. (A) Periosteal stem cell (hPSC), periosteal progenitor 1 (hPP1), and periosteal progenitor 2 (hPP2) populations described in (12) overlaid onto viSNE plots of different tissues. (B-E) Gating to identify hPSCs, hPP1, and hPP2 in (B) periosteum, (C) cartilage (D), endosteum, and (E) bone marrow. (F-H) Frequency of (F) hPSC, (G) hPP1, and (H) hPP2 on Lin-fractions of periosteum, cartilage, endosteum, and bone marrow samples, n=21. *p<0.05, **p<0.01, ***p<0.001, ****p<0.0001, one-way ANOVA with Turkey’s post hoc test.

**Figure 4-figure supplement 1.**
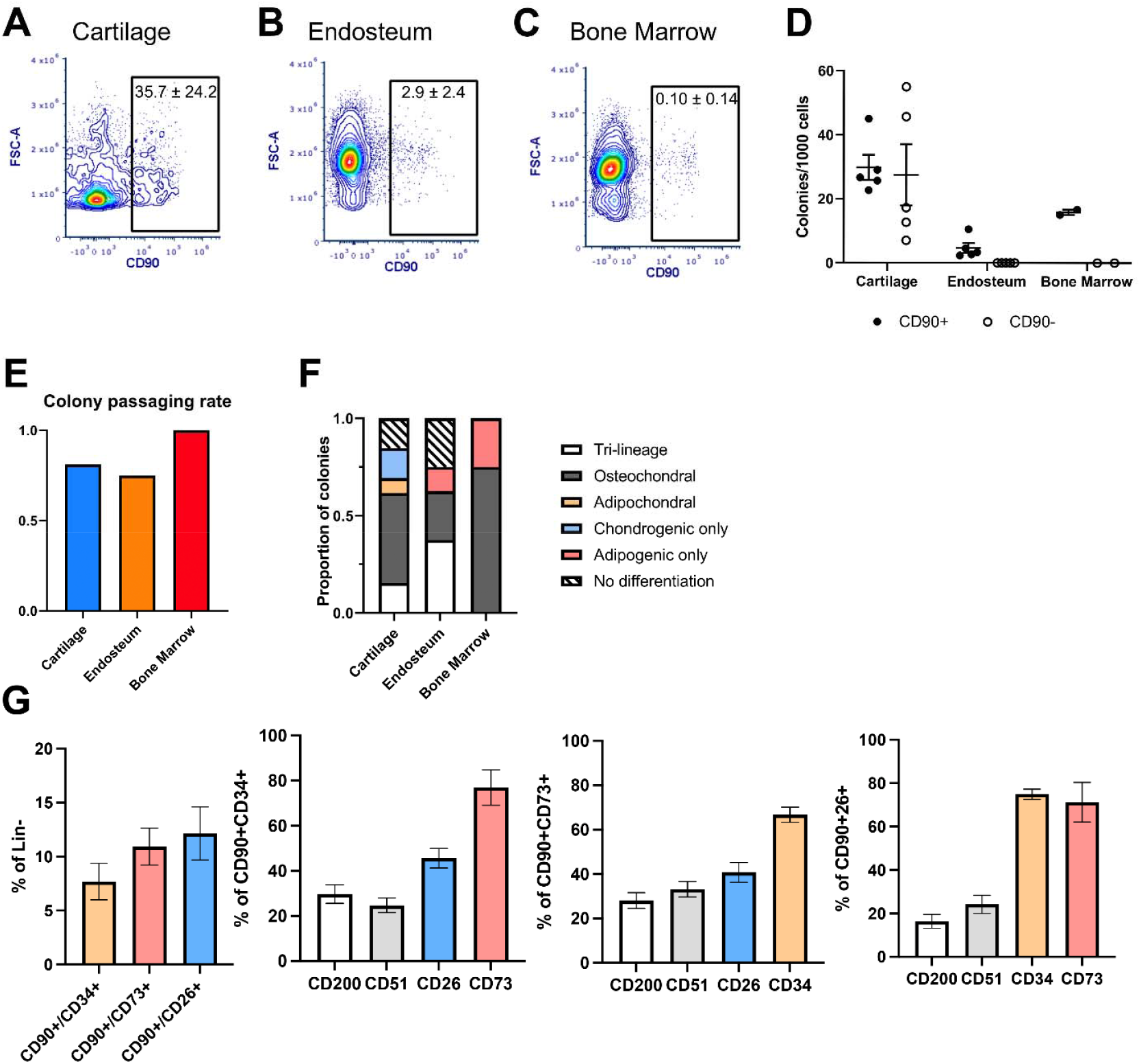
Growth and differentiation potential of CD90+ cells from cartilage, endosteum and bone marrow. Representative density plot of CD90 expression in the (A) cartilage, (B) endosteum, and (C) bone marrow, the percentages indicate the proportion of events ± standard deviation from sorting experiments, n=2-5. (D) Comparison of colony-forming unit fibroblast (CFU-F) frequency in CD90- and CD90+ populations. (E) Passaging rate of the single colonies from CD90+, four colonies are picked from each tissue, n=1-5. (F) Differentiation potential of expanded CD90+ colonies. (G) Expression of selected markers within the indicated CD90+ subpopulations in the periosteum that were evaluated ex vivo in Figure 4, n=16-18.

**Figure 2-figure supplement 2.**
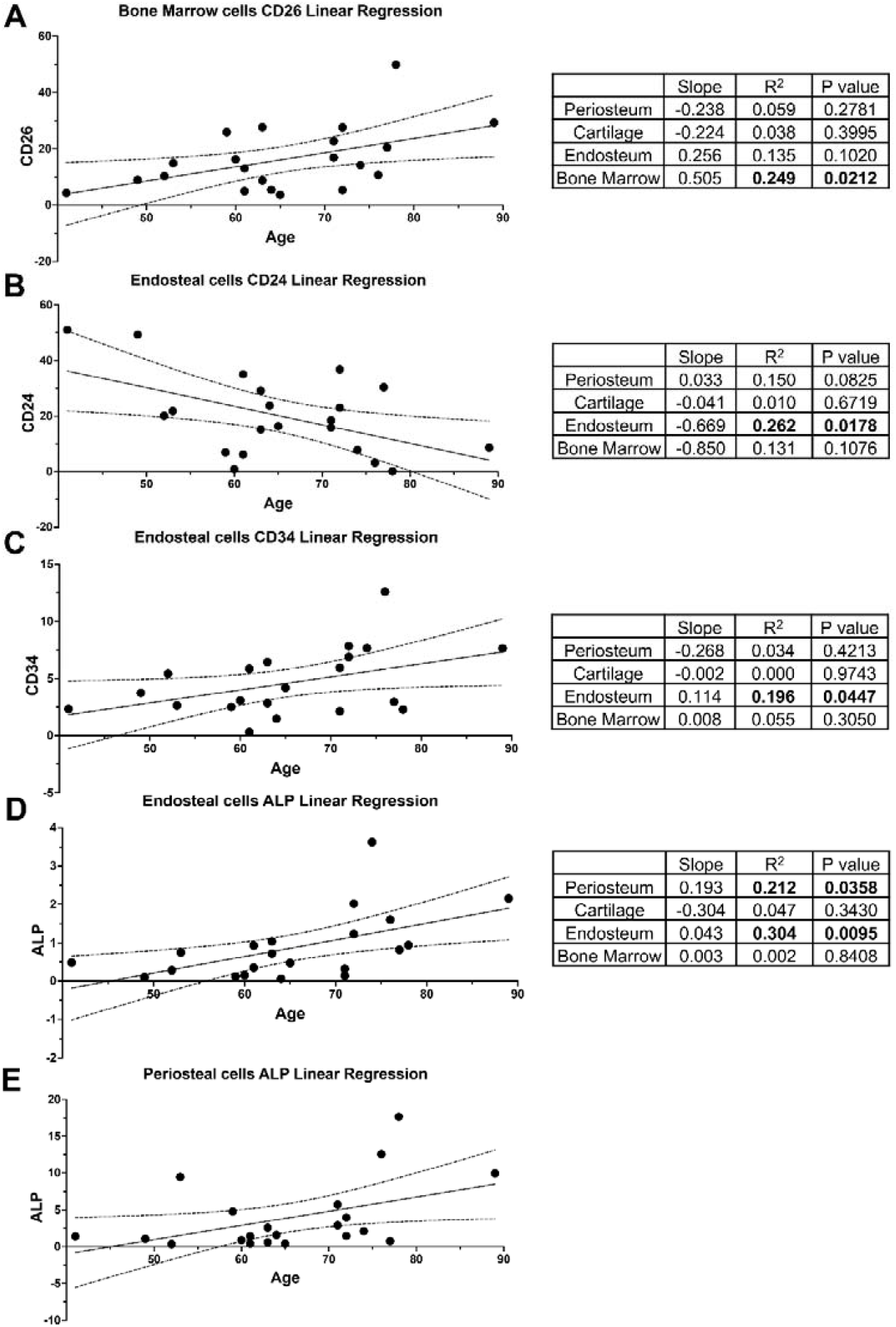
Markers that were affected by age. Simple linear regression analysis of each individual marker with age were performed for individual tissues. Correlations where the slope was significantly non-zero are shown. Data are shown with 95% confidence bands of the best-fit line, n=21. The slope, R^2^, and p-value for all tissues for each marker shown are indicated in the tables on the right.

## Additional legends

Figure 1 – Source data 1: Raw data for Figure 1C and Figure supplements

Figure 2 – Source data 1: Raw data and full statistical analysis for Figure 2 and Figure supplement 2

Figure 2 – Source data 2: Raw data for Figure 2 supplement 1, analysis of Lin+ populations

Figure 3 – Source data 1: Raw data and full statistical analysis for Figure 3 and supplement

Figure 4 – Source data 1: Raw data and full statistical analysis for Figure 4 and supplement

Figure 5 – Source data 1: Raw data and full statistical analysis for Figure 5

